# Landscape of tumor-infiltrating immune cells in human immunogenic cancers:B cells are inversely associated with CD8T cells, but positively correlated with Treg cells

**DOI:** 10.1101/641712

**Authors:** Young Kwang Chae, William Han Bae, Minji Jung, Young Suk Kim, Jonathan Forrest Anker, Keerthi Tamragouri, Maria Matsangou, Francis Joseph Giles

## Abstract

The composition of tumor-infiltrating immune cells may be a strong predictor of cancer treatment responses and survival outcomes. While B cells have been suggested to suppress T cell cytotoxicity in preclinical studies, it has been less understood whether B cells will exert immune-regulatory roles in human cancers. We explored immune cell landscapes in six human immunogenic cancers, including bladder cancer, head and neck cancer, lung adenocarcinoma, lung squamous cell carcinoma, melanoma, and renal cell carcinoma by calculating gene expression patterns of immune cell-specific metagenes in a total of 2951 cancers. We demonstrated that tumor-infiltrating activated B cells was correlated with regulatory T cell (Treg) infiltration, but had an inverse association with activated CD8 T cell infiltration consistently across all six human cancers. Tumors infiltrated by activated B cells (ActB+ tumors) demonstrated an elevated expression of regulatory cytokines and immune checkpoints, compared to tumors without infiltration by activated B cells (ActB-tumors). Activated B infiltration was not significantly associated with survival outcomes.

**Précis:** This human cancer tissue analysis showed that tumor infiltration by activated B cells correlates with decreased infiltration by activated CD8 T cells in immunogenic solid tumors, implicating B cell inhibition may enhance T cell-mediated cytotoxicity.

## Introduction

Immunotherapy has become an attractive cancer treatment option in clinical practice. This new therapeutic approach has yielded favorable survival outcomes with fewer treatment-related adverse events compared to conventional chemotherapeutic regimens in multiple human cancers [1–3]. These agents target immune checkpoint molecules, such as programmed cell death protein 1 (PD-1) and cytotoxic T-lymphocyte-associated protein 4 (CTLA-4), to block the inhibition of T lymphocyte activity. The Food and Drug Administration (FDA) approved the use of immune checkpoint inhibitors in many solid tumors, including melanoma, renal cell carcinoma, head and neck cancer, non-small cell lung cancer, and bladder cancer. However, accumulating data suggest that only a small proportion of patients responds to this treatment approach [4,5], and many responders go on to develop acquired drug resistance [6,7]. It is not yet well understood what causes differential responses among patients and how many initially responding tumors eventually are able to evade host anti-tumor immunity.

One mechanism of tumor immunoevasion is the T cell suppression induced by other tumor-infiltrating immune cells, such as regulatory T cells (Tregs) and myeloid-derived suppressor cells (MDSCs). Tregs downregulate CD8 T cell proliferation and cytolytic activity in multiple cancer types [8–10]. In fact, decreased tumor-infiltrating CD8 T cells to Tregs ratio is a strong predictor of poor prognostic outcomes in ovarian, gastric, and breast cancers [11–14]. MDSCs are also implicated in T cell suppression [15], tumor progression [16], poor survival outcomes [17], and increased resistance to anti-cancer therapies [18,19]. Despite recent progress in our understanding of the pro-tumorigenic roles of Tregs and MDSCs, less is known about the immunosuppressive properties of tumor-infiltrating B cells.

Recent preclinical studies have suggested that tumor-infiltrating B cells may be directly or indirectly responsible for the T cell inactivation and disease progression. Qin *et al.* reported that mammary adenocarcinoma tissues implanted to the B cell depleted mouse (BCDM) models show no growth, in comparison to tumor development in almost all wildtype (WT) mice [20]. The decreased tumor development in BCDM models was associated with enhanced CD8 T cell cytotoxic activities. Another study using the BCDM models with thymoma and colorectal cancer tissue implants also reported spontaneous tumor regression, higher Th1 cytokine responses, and increased CTL responses, all of which were reversed upon adoptive transfer of B cells [21]. Consistent with these findings, combination treatment with anti-CD20 monoclonal antibodies and platinum- or taxol-based chemotherapeutics resulted in faster tumor regression and increased CD8 T cell infiltration than chemotherapy alone in murine squamous cell carcinoma [22]. The enhanced CD8 T cytotoxicity associated with the B cell depletion suggest that B cells are a key element in tumor microenvironment(TME) immunosuppression.

To explain this B cell-mediated T cell suppression, many researchers have focused on the role of regulatory B cells (Bregs), a small subpopulation of B cells characterized by their ability to suppress inflammatory responses. Tumor-associated stimuli appear to potentiate the differentiation of B cells into Breg cells to suppress T cell-mediated immune responses. *In vitro* co-culture of murine splenic B cells with CD4 and CD8 T cells in 4T1 breast cancer cell-derived media (CM) resulted in decreased T cell proliferation [23]. These tumor-evoked B cells that phenotypically resembled activated mature B cells constitutively expressed Stat3 and induced conversion of resting CD4 T cells to Foxp3+ Tregs. This study implicated that B cells may acquire a new immune-regulatory property after exposure to cancer-associated signals, leading to immune tolerance in the TME.

Even though preclinical studies provide evidence that tumor-infiltrating B cells may directly or indirectly affect anti-tumor T cell responses in the TME, the interaction between B and T cells has not been thoroughly evaluated in human cancer histologies. We analyzed expression levels of immune “metagene” markers specific for 28 distinct immune cell types to predict tumor infiltration in six selected human cancers that has FDA indication for T-cell medicated immune checkpoint therapy: bladder cancer, head and neck cancer (HNC), lung squamous cell carcinoma (lung SCC), lung adenocarcinoma, melanoma, and renal cell carcinoma (RCC). Further, infiltration patterns of immune cells were analyzed in relation to the presence of tumor-infiltrating B cells. Here, we report that the tumor-infiltrating B cells are inversely associated with activated CD8 T cells in all six human immunogenic cancers.

## Materials and Methods

This study utilized cBioPortal [24,25] to obtain the RNA-seq values and clinical data of 2951 patients with bladder urothelial carcinoma, head and neck squamous cell carcinoma, lung adenocarcinoma, lung SCC, cutaneous melanoma, and RCC from The Cancer Genome Atlas (TCGA) Research Network [26]. The mRNA expression z-scores of the previously published 812 immune metagenes were analyzed using the Gene Set Enrichment Analysis (GSEA) software from the Broad Institute to predict the tumor infiltration by 28 distinct immune cell types [27–29]. Any immune cell types with a false discovery rate (q-values) less 10% were considered positively infiltrating [29].

We calculated odds ratios (ORs) of the number of tumors with versus without infiltration by each respective immune cell type in respect to the presence of activated CD8T cells according to the following equation for all six histologies. ORs for activated B cells were also calculated in the same manner.

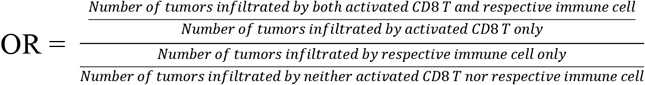

## Results

### Six human immunogenic cancers were most infiltrated by activated CD4 T and activated CD8 T cells

To understand the landscapes of infiltrating immune cells in human cancers, we analyzed expression patterns of “immune metagenes” to predict infiltration by 28 distinct immune cell types in 2951 tumors across six cancer types. 1751 (59.33 %) and 253 (8.57 %) samples were infiltrated by any T and B cell types, respectively (Fig. 1). Human cancers were most frequently infiltrated by activated CD8 (27.34%) and activated CD4 (21.35%) T cells. Infiltration by activated and immature B cells were identified in 165 (5.59%), and 111 (3.76%) tumors, respectively. Additionally, MDSCs and Treg cells were present in 37 (1.25%) and 67 (2.27%) tumors, respectively. Previously we demonstrated that activated CD8 and CD4 T cells were the most common immune cell types within non-small cell lung cancer TMEs [30,31]. Similar immune cell landscapes were observed in our prediction models for other four human cancers (Supplementary Fig. S1A-S1D).

**Figure 1.**
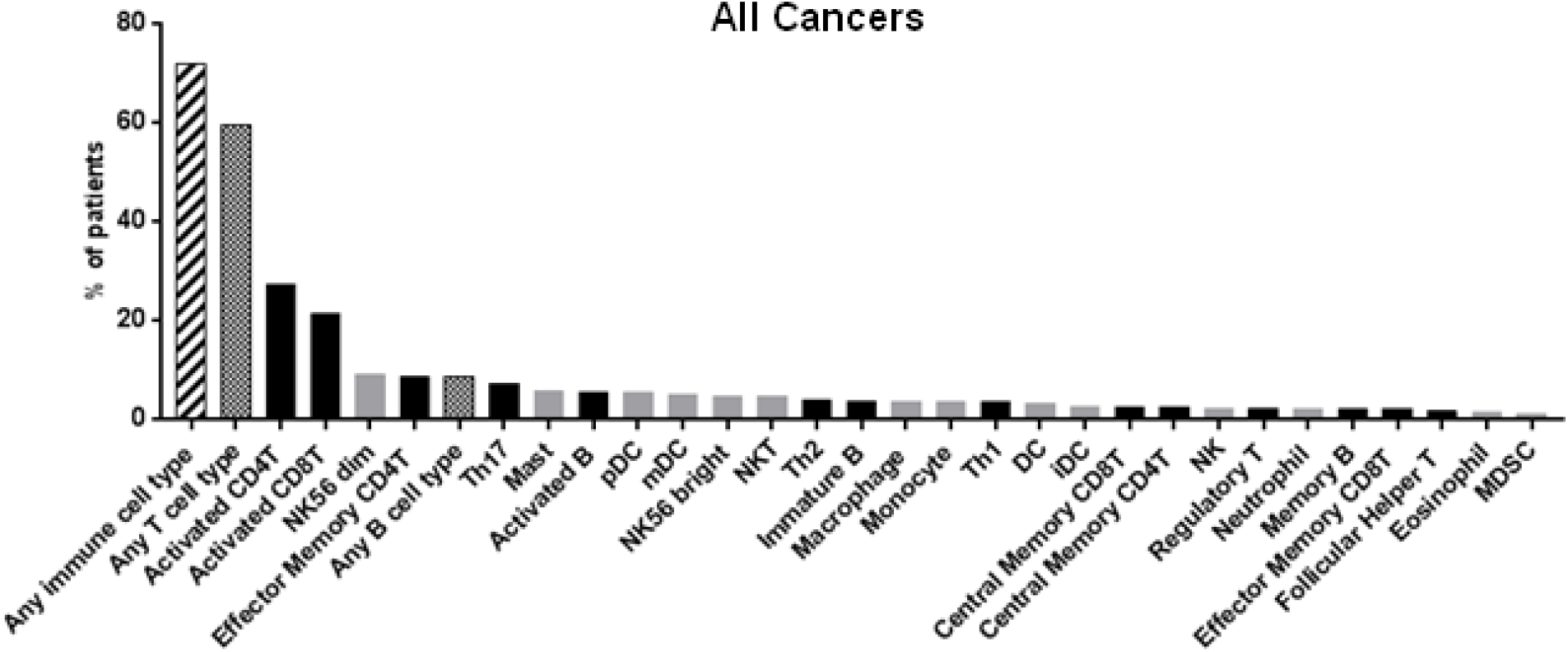
Percent of tumors infiltrated by each immune cell type in all immunogenic cancers. NK56 bright, CD56 bright natural killer cell; DC, dendritic cell; iDC, immature dendritic cell; pDC, plasmacytoid dendritic cell; mDC, mature dendritic cell; NKT, natural killer T cells; MDSC, myeloid-derived suppressor cells (Bold bars = adaptive immune cell types, gray bars = innate immune cell types).

### Infiltration by activated CD8 T cells occurred differentially in tumors with activated B cell infiltration compared to ones without

TMEs appear to modulate the functional orientation of cytotoxic T cells within tumor tissues. Tumor cells and other infiltrating immune cells express or secret immuno-modulating mediators, such as cytokines, chemokines, and immune checkpoints, to regulate the CD8 T cell infiltration and their cytotoxic activities. To evaluate how infiltration by other immune cell types may impact the presence of activated CD8 T cells, we calculated the ORs of the numbers of tumors with versus without infiltration by each immune cells with respect to the presence of activated CD8 T cells in the six cancer histologies individually and in all cancers combined (All Cancers). The All Cancers group exhibited a significant association between activated CD8 T cell infiltration and infiltration by CD56 bright natural killer cells (NK56 bright), mature dendritic cells (mDC), effector memory CD4 T cells, activated CD4 T cells, CD56 dim natural killer cells (NK56 dim), Th17 cells, memory B cells, and monocytes. Further, tumors with infiltration by activated CD8 T cells were significantly less likely to contain infiltration of Th2 cells, macrophages, dendritic cells (DC), effector memory CD8 T cells, Th1 cells, activated B cells, eosinophils, Tregs, central memory CD4 T cells, central memory CD8 T cells, immature B cells, and NK cells (Fig. 2A).

**Figure 2.**
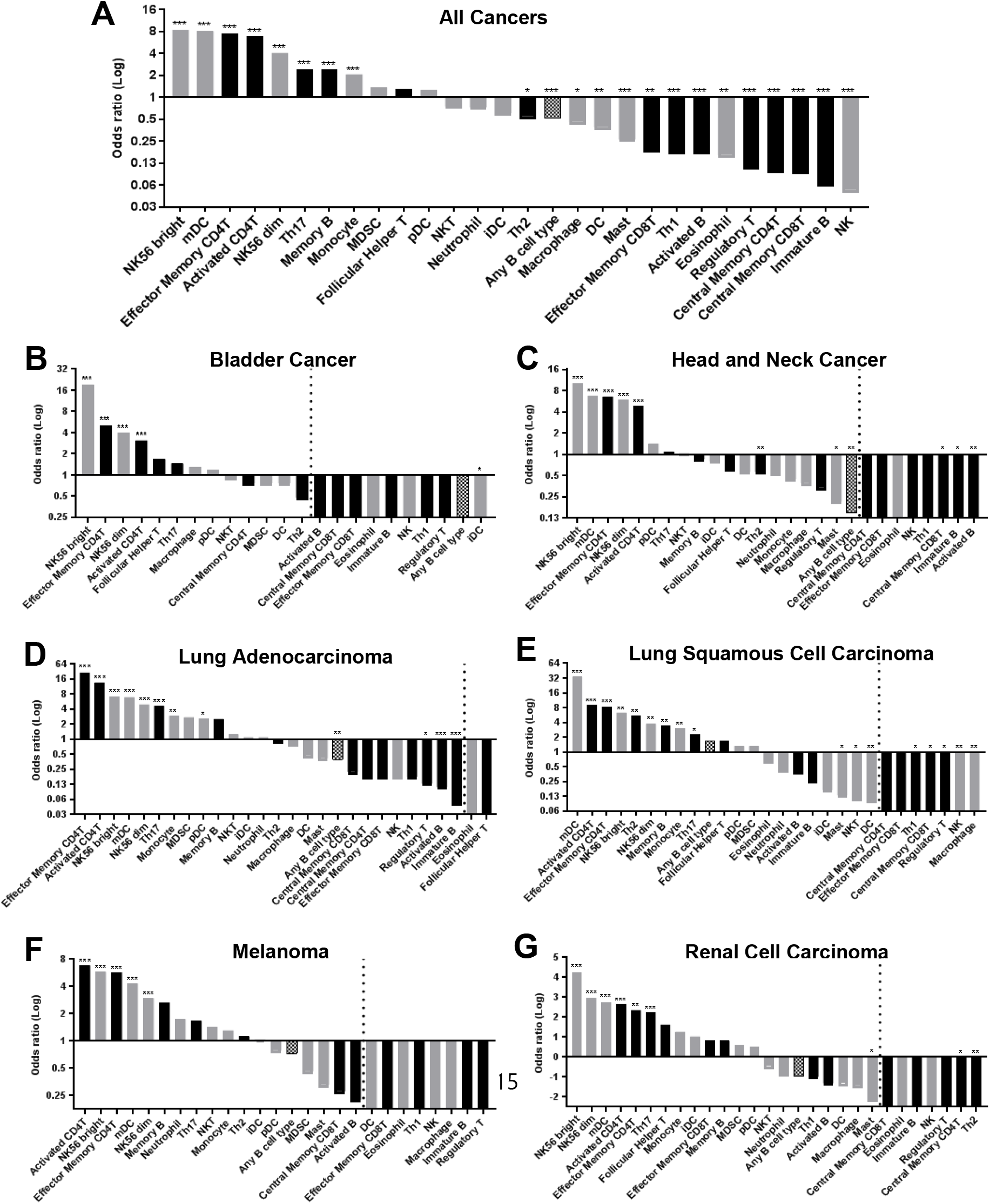
Odds ratios of the number of tumors with versus without infiltration by each immune cell with respect to the presence of activated CD8T cells. Located from the top represent the odds ratios in (**A**) all cancers, (**B**) bladder cancer, (**C**) head and neck cancer (**D**) lung adenocarcinoma, (**E**) lung squamous cell carcinoma, (**F**) melanoma, and (**G**) renal cell carcinoma. Cell types on the right of the dotted lines were never co-infiltrated by activated CD8T cells (Immune cell types with tumor infiltration in less than three samples were excluded). *** *P*<0.001, ** *P*<0.01, * *P*<0.05

Individually in all six cancer histologies, co-infiltration was found between activated CD8 T cells and activated CD4 T cells, effector memory CD4 T cells, NK56 bright cells, and NK56 dim cells (Fig. 2B-2G). Mature DCs were significantly present in the same tumors with activated CD8 T cells in all histologies except bladder cancer. Th17 cells were significantly infiltrated in the same tumors with activated CD8 T cells in lung adenocarcinoma, lung SCC, and RCC (Fig. 2D, 2E, and 2G). Monocyte and pDC infiltrated tumors with activated CD8 T cells in lung adenocarcinoma (Fig. 2D). Th2 cells, memory B cells, and monocyte co-infiltrated tumors with CD8 T cells in lung SCC (Fig. 2E).

In bladder cancer, iDC was significantly infiltrated in other tumors than activated CD8 T cells (Fig. 2B). In HNC, any B cell type, activated B cells, immature B cells, Th2 cells, mast cells, and central memory CD8 T cells were significantly inversely associated with activated CD8 T cells (Fig. 2C). Similarly, in lung adenocarcinoma, infiltration by any B cell type, activated B cells, immature B cells, and Tregs was significantly mutually exclusive with infiltration by activated CD8 T cells (Fig. 2D). In lung SCC, mast cells, NKT, DCs, Th1 cells, central memory CD8 T cells, Tregs, NK cells, and macrophages had significant inverse patterns of tumor infiltration with activated CD8 T cells (Fig. 2E). In RCC, Th2, central memory CD4T, and mast cells were significantly inversely associated with activated CD8 T cells (Fig. 2G). Interestingly, infiltration by activated and immature B cells demonstrated consistent inverse associations with infiltration by activated CD8 T cells across all six human cancers (Fig. 2B-2G; ORs<1).

### Activated B cell infiltration was associated with Treg infiltration in all six human cancers

To further investigate how B cell infiltration may affect the immunophenotype in human cancer TMEs, we calculated the ORs of the numbers of tumors with versus without infiltration by each immune cell with respect to the presence of activated B cells.

In All Cancers, activated B cells were significantly co-infiltrated in the same tumors with Tregs, macrophages, immature B cells, effector memory CD8 T cells, central memory CD8 T cells, central memory CD4 T cells, Th1 cells, NK cells, and any T cells. On the contrary, there was significant mutual exclusivity between infiltration by activated B cells and infiltration by activated CD8 T cells, activated CD4 T cells, effector memory CD4 T cells, Th17 cells, Th2 cells, NK56 dim cells, mature DCs, plasmacytoid DCs, and monocytes (Fig. 3A). Notably, Activated B cells significantly co-infiltrated tumors with Tregs in All Cancers and five individual cancers except melanoma (Fig. 3). Tregs had a non-significant association with activated B cells in melanoma (OR=5.05, p=0.11; Fig. 3F). Consistently, Treg-specific markers *Foxp3*, *CD25*, and *CTLA4* were significantly elevated in tumors with (ActB+ tumors) versus without (ActB- tumors; p<0.001; Fig. 4B) infiltration by activated B cells.

**Figure 3.**
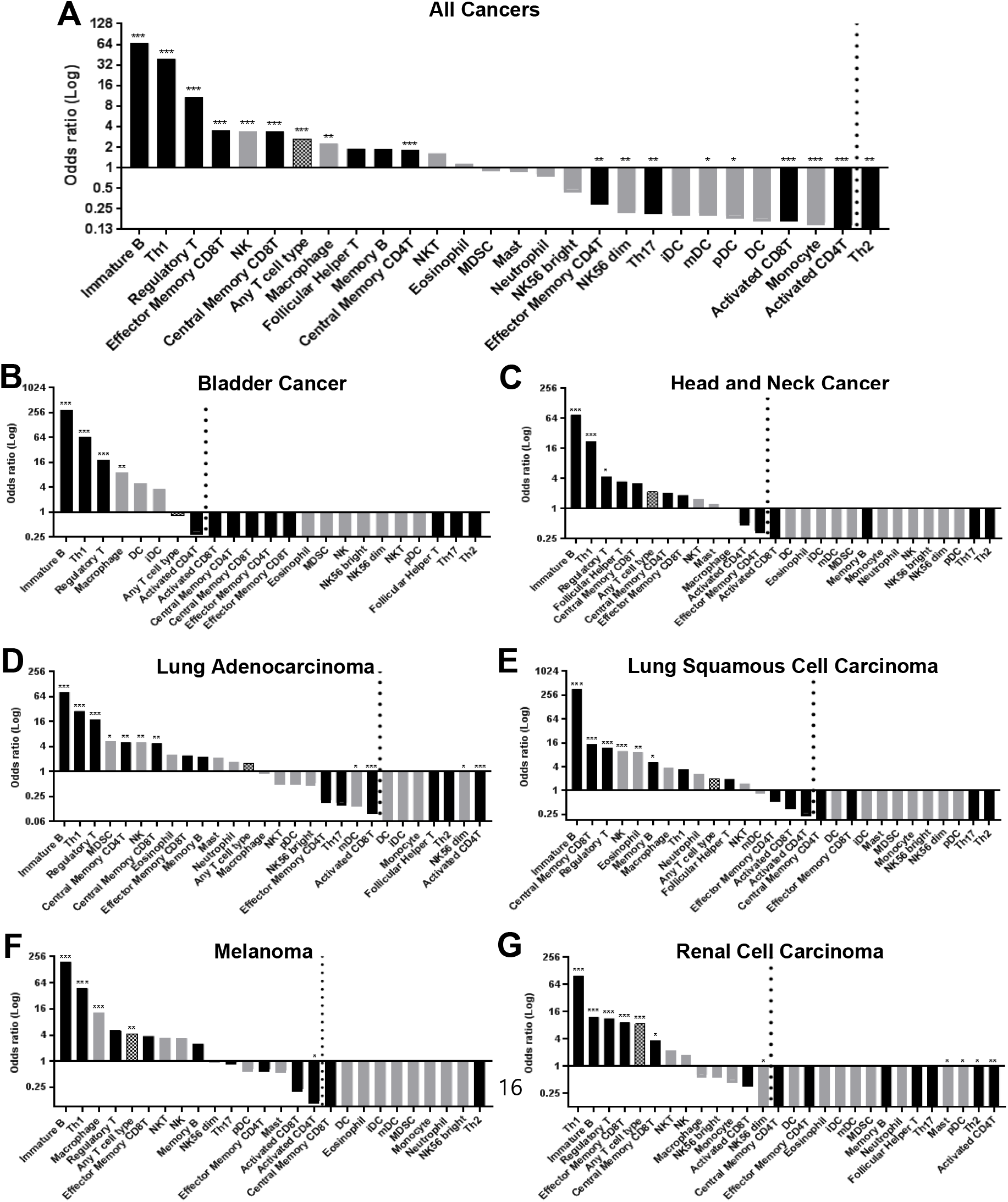
Odds ratios of the number of tumors with versus without infiltration by each immune cell with respect to the presence of activated B cells. Located from the top represent the odds ratios in (**A**) all cancers, (**B**) bladder cancer, (**C**) head and neck cancer (**D**) lung adenocarcinoma, (**E**) lung squamous cell carcinoma, (**F**) melanoma, and (**G**) renal cell carcinoma. Cell types on the right of the dotted lines were never co-infiltrated by activated B cells (Immune cell types with tumor infiltration in less than three samples were excluded).

Activated B cells significantly co-infiltrated tumors with immature B cells in all six histologies (Fig. 3B-3G), with Th1 cells in all cancers except lung SCC (Fig. 3B-3G), with macrophages in bladder cancer and melanoma (Fig. 3B and 3F),with MDSCs, central memory CD4 T cells, central memory CD8 T cells, and NK cells in lung adenocarcinoma (Fig. 3D), with memory B cells, eosinophils, NK cells, and central memory CD8 T cells in lung SCC (Fig. 3E), and with any T cells, effector memory CD8 T cells and central memory CD8 T cells in RCC (Fig. 3G).

### Tumor tissues infiltrated by activated B cell were associated with higher expressions of regulatory cytokines, immune checkpoints, and Fcγ receptors (FcγRs)

Because expansion of Breg cells has been associated with T cell suppression in preclinical cancer models, we analyzed expression of Breg-associated markers in ActB+ and ActB- tumors. Regulatory cytokines *IL10* and *IL35* (*Ebi* and *12p35*) were expressed significantly higher in ActB+ than ActB- tumors, while cytokine *TGFβ1* showed no change between tumor stratifications (Fig. 4A). Transcription factor *Stat3*, which is constitutively expressed in tumor-evoked Bregs [23] and immune checkpoint *PD1* was also expressed significantly higher in ActB+ tumors(Fig. 4B-C).

**Figure 4.**
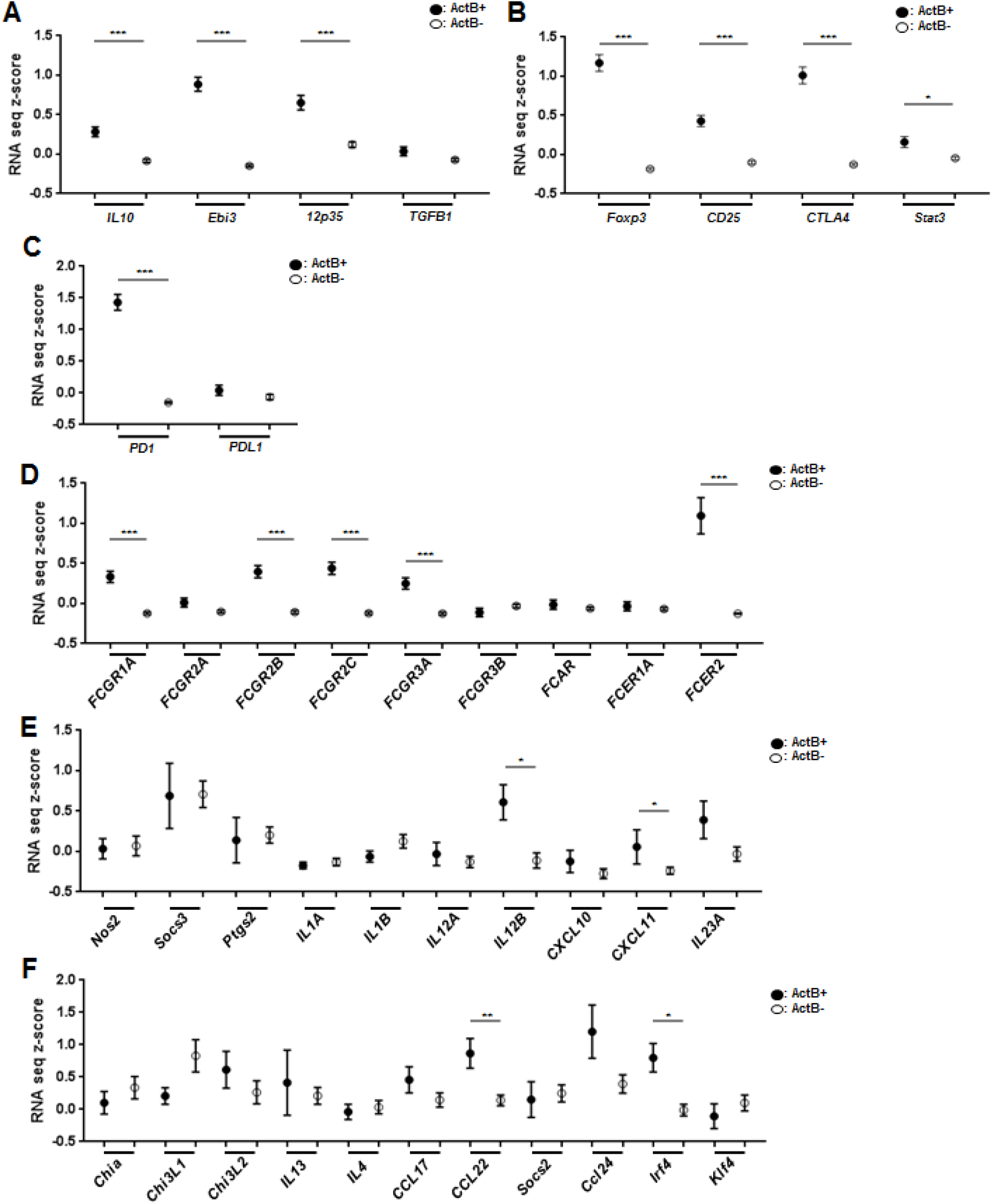
Expression scores in tumors with activated B cell infiltration (ActB+) and tumors without (ActB-): (**A**) Regulatory cytokines IL10 (*IL10*), IL35 (*Ebi3, 12p35*), and TGFβ(*TGFB1*), (**B**) regulatory cell-associated markers *Foxp3, CD25, CTLA4*, and *Stat3*, (**C**) immune checkpoints *PD1* and *PDL1*, and (**D**) Fcγ receptors. (**E-F**) M1-phenotype (E) and M2-phenotype associated markers (F) in tumors infiltrated by macrophages

Previously, it has been established that activation of FcγRs drives carcinogenesis and tumor progression in murine models [32]. *FCGR1A*, *FCGR2B*, *FCGR2C*, and *FCGR3A* expression were significantly elevated in ActB+ tumors (Fig. 4D). While immune complex binding to FcγRs on macrophages has been reported to repolarize these cells toward an M2 pro-tumorigenic phenotype [32], we did not observe a significant association between activated B cell infiltration and elevated expression of M1(Fig. 4E) or M2 polarization markers(Fig. 4F) in tumor tissues infiltrated by macrophages.

### Expressions of cytotoxicity-associated markers and anti-angiogenic markers were elevated in ActB+ tumors

Despite the inverse association between activated B and CD8 T cells in human cancers (Fig. 2), cytotoxicity-associated markers *Prf1, Gzma, Gzmb*, and *Ifng* were significantly higher in ActB+ tumors (Supplementary Fig. S2A). Previous histological analyses revealed the presence of lymphoid structures within tumor tissues, called Tertiary Lymphoid Structure (TLS), which is characterized by clusters of mature DC, T cells and follicular B cells, has been associated with Th1 polarization and increased CD8 T toxicity [33]. Consistent with the finding, expression score of TLS markers CXCL13 and IL21 were significantly increased in ActB+ tumors (Supplementary Fig. S2B). Furthermore, CXCL13 expression score among ActB+ tumors had positive correlation with *Gzma* and *Gzmb* (Supplementary Fig. S2C-D; r=0.179, p<0.05; r=0.153, p=0.05). No association was found between CXCL13 and *Prf1* (Supplementary Fig. S2E).

Anti-angiogenic makers, *Cxcl4*, *Cxcl9*, *Cxcl10*, *Cxcl11*, and *Cxcr3* were elevated in the presence of infiltrating activated B cells while angiogenic markers showed differential expression patterns (Supplementary Fig. S3).

### Infiltration by activated B cells was not associated with patient survival outcomes

B cell infiltration was not significantly associated with overall survivals in all human cancers except HNC (Supplementary Fig. S4). Median overall survival was 16.8 months longer in HNC ActB+ tumors (p<0.05; Supplementary Fig. S4B). Disease-free survivals also showed no significant association with B cell infiltration in all six cancer histologies (Supplementary Fig. S5).

## Discussion

Consistent with preclinical findings that B and T cells are functionally counteractive, here in this human cancer tissue analysis, we identified that infiltration by activated B cells was inversely related to infiltration by activated CD8 T cells. Although previous functional studies all have suggested the inverse relationship, this has not been shown at the human tissue level.

We analyzed expression scores of the immune metagenes to predict immune cell landscapes in six human cancers [29]. More than half of tumors were infiltrated by any T cell types, and approximately one in twelve tumors were infiltrated by any B cell types. Activated CD4 T and CD8 T cells were most common immune cell types, present in roughly one in four and one in five human cancers, respectively. ActB+ tumors were less frequently infiltrated by activated CD8 T cells compared to ActB- tumors (p<0.001). This mutually exclusive infiltration pattern between activated B and CD8 T cells was observed consistently across all six human cancers investigated.

Even though B cells activate the immune system via antigen presentation and antibody production, they also exhibit immunosuppressive functions in chronic inflammatory conditions. Breg cells inhibit pro-inflammatory immune responses primarily via the production of regulatory cytokines, such as IL10 and IL35. IL10 inhibits T cell proliferation [34], potentiates Treg cell differentiation [35], and enhances expression of CTLA4 in Foxp3+ Treg cells via Stat3 signaling pathway [35]. Due to the universality of IL10 expression by multiple Breg subpopulations, this cytokine is commonly utilized as an identifier of Breg cells. Similarly, IL35, another major regulatory cytokine produced by Breg cells, also prohibits T cell proliferation and anti-tumor functions while promoting proliferation of Tregs [36]. Consistently in our human cancer tissue analysis, ActB+tumors showed a significantly higher expression score of regulatory cytokines IL10 and IL35 compared to ActB- tumors (Fig 4A). The elevation of the unestablished Breg markers in ActB+ tumor, coinciding with suppression of CD8 T cells, appear to suggest increased infiltration by Breg cells. However, the clear association between Breg and Act CD8 T cells could not be demonstrated in this study due to the lack of specific markers for Breg cells

In accordance with the elevated expression of regulatory cytokines in ActB+ tumors, we observed a positive correlation between tumor-infiltrating activated B cells and Tregs in the All Cancers group and multiple individual tumor types (Fig. 3). A significant elevation of Treg-specific markers *Foxp3*, *CD25*, and *CTLA4* were identified in ActB+ tumors (p<0.001; Fig. 4A). These findings are consistent with the *in vitro* finding that cancer-associated stimuli “educate” B cells to induce expansion of Treg cells to indirectly inhibit proliferation of CD8 T cells [23]. An increased expression of immune checkpoint *PD-1* was also noted in the presence of infiltrating activated B cells (p<0.001 for all three; Fig. 4C). Given the recent isolation of IL10-producing PD1^high^ Breg cells in human hepatocellular carcinoma [37], it is plausible to suspect that PD1^high^ Breg cell may also exist in other human cancer types. Or, because activated B cells and Treg cells were significantly present in the same tumors, the elevated expression of immune checkpoints in ActB+ tumors may be largely attributed to infiltrating Treg cells.

In lung adenocarcinoma, tumor-infiltrating activated B cells were significantly associated with infiltrating MDSCs (p<0.05; Fig. 3D). MDSCs may lead to T cell suppression via the provision of reactive nitrogen and oxygen species, such as NO, ROS, and H_2_O_2_ [38]. Our finding is consistent with previous analyses of peripheral blood of NSCLC patients that found a significant correlation between IL-10 producing-Breg cells and MDSCs [39]. Other cancers did not have a significant association between B cells and MDSCs. However, it is noteworthy to highlight that MDSCs may exert a stronger regulatory function to suppress T cell proliferation in the presence of B cells. In a previous preclinical study, while implantation of syngeneic mammary cancer tissues led to a comparable expansion of MDSCs in both BCDM and WT mice, MDSCs from WT mice exhibited an increased production of ROS and H_2_O_2_, and more efficiently suppressed of T cell proliferation than MDSCs from BCDMs [40]. However, we did not observe a significant elevation of nitric oxide synthase (iNOS) and arginase 1 (Arg1) associated with co-infiltration by activated B cells and MDSCs, possibly due to the small sample size (n=2; data not shown).

We found that ActB+ tumors contained significantly elevated FcγRs expression and increased macrophage infiltration (Fig. 3A, 4D). Activation of FcγRs causes robust angiogenic responses and recruitment of pro-tumorigenic leukocytes [32]. Additionally, immune complex accumulation in the neoplastic tissues is known to stimulate FcγRs to repolarize macrophages into tumor-promoting M2 phenotypes. However, we did not observe a significant association between tumor-infiltrating B cells, expression of angiogenic markers, and M2 macrophage polarization in the tumor tissue levels (Fig. 4E, 4F). This finding is in line with the previously suggested notion that peripheral B cells may play more important roles in providing tumor-infiltrating immune complexes via distal production of antibody, and therefore determines polarization status of tumor-infiltrating macrophages [32]. Due to the inherent limitation of our human tissue analysis, we were unable to isolate macrophages to evaluate their functional phenotypes at the cellular level. Our finding does not exclude that the tumor-infiltrating B cells may influence the macrophage phenotypes in human cancer microenvironment.

Interestingly, activated B cell infiltration was associated with elevated expression of T cell cytotoxicity-associated markers *Gzma, Gzmb, Prf1* and *Ifng* (Supplementary Fig. S2A) and infiltration by Th1 cells (Fig. 3). These findings may be attributed to the functional diversity of B cells within tumor tissues. Previous histopathological analyses revealed that B cells are mostly located in either tertiary lymphoid structures (TLS) or invasive margin of growing tumors [33]. TLS, as an active site of inflammatory responses, comprises of germinal centers where proliferating follicular B cells and helper T cells reside, and surrounding T cell zone where naive T cells mature into effector T cells with the help of mature DCs. The presence of TLS has been linked to strong Th1 responses, enhanced antibody production, and superior survival outcomes in NSCLCs [41,42]. The correlation between activated B cell and Th1 responses (Supplementary Fig. S2A) may be attributed to the co-localization of B cells and CD8 T cells in the active areas of inflammation. Consistently, TLS-associated marker *CXCL13* was a strong correlate of *Gzma* and *Gzmb* in ActB+ tumors (S1C-E). However, it is not clear how frequently TLS formations are found within solid tumors and how B cells in invasive margins behave differently from ones in TLS. Due to the inherent limitation of our experimental design, we were unable to compare the functional orientations of these two B cell groups.

Preclinical studies provide evidence that inhibition of B cells enhances T cell-mediated antitumor immunity, leading to tumor regression and favorable survival outcomes. A combination treatment regimen of Bruton’s Tyrosine Kinase (BTK) inhibitor and PD-L1 inhibitor in murine colon and breast cancer models led to enhanced T cell cytotoxic activities and prolonged survivals compared to the mice treated with PD-L1 and BTK inhibitor monotherapies [43]. Another study on pancreatic ductal adenocarcinoma demonstrated that addition of a BTK inhibitor with gemcitabine increased the CD8 T cell infiltration and T cell cytotoxic responses in mice, resulting in tumor growth retardation [44]. Similarly, administration of anti-CD20 monoclonal antibody in K14-HPV16 head and neck squamous carcinoma mouse resulted in increased infiltration of CD8 T cells, which in turn may have contributed to the decreased tumor growth and increased sensitivity to cytotoxic agents [22]. Further, analysis of peripheral blood of human NSCLC patients revealed a significant association between the frequency of IL-10 producing B cells and increased histopathological stage [39]. Based on these findings, our group has designed a clinical trial using a combination therapy of ublituximab, an anti-CD20 monoclonal antibody, and nivolumab, an anti-PD-1 inhibitor, for the treatment of NSCLC and HNSCC patients, which will be opening soon.

In summary, while preclinical findings demonstrated regulatory roles of B cells to impede T cell anti-tumor activity, to date, there has been no validating data with human tissues. We utilized RNA expression scores for immune-related metagenes to predict immune cell landscapes in six human cancers. Furthermore, we demonstrated, for the first time, that tumor infiltration by B cell was inversely associated with infiltration by activated CD8 T cells, while positive correlating with infiltration by Tregs in multiple human cancers.

---

**Supplementary Figure 1.**
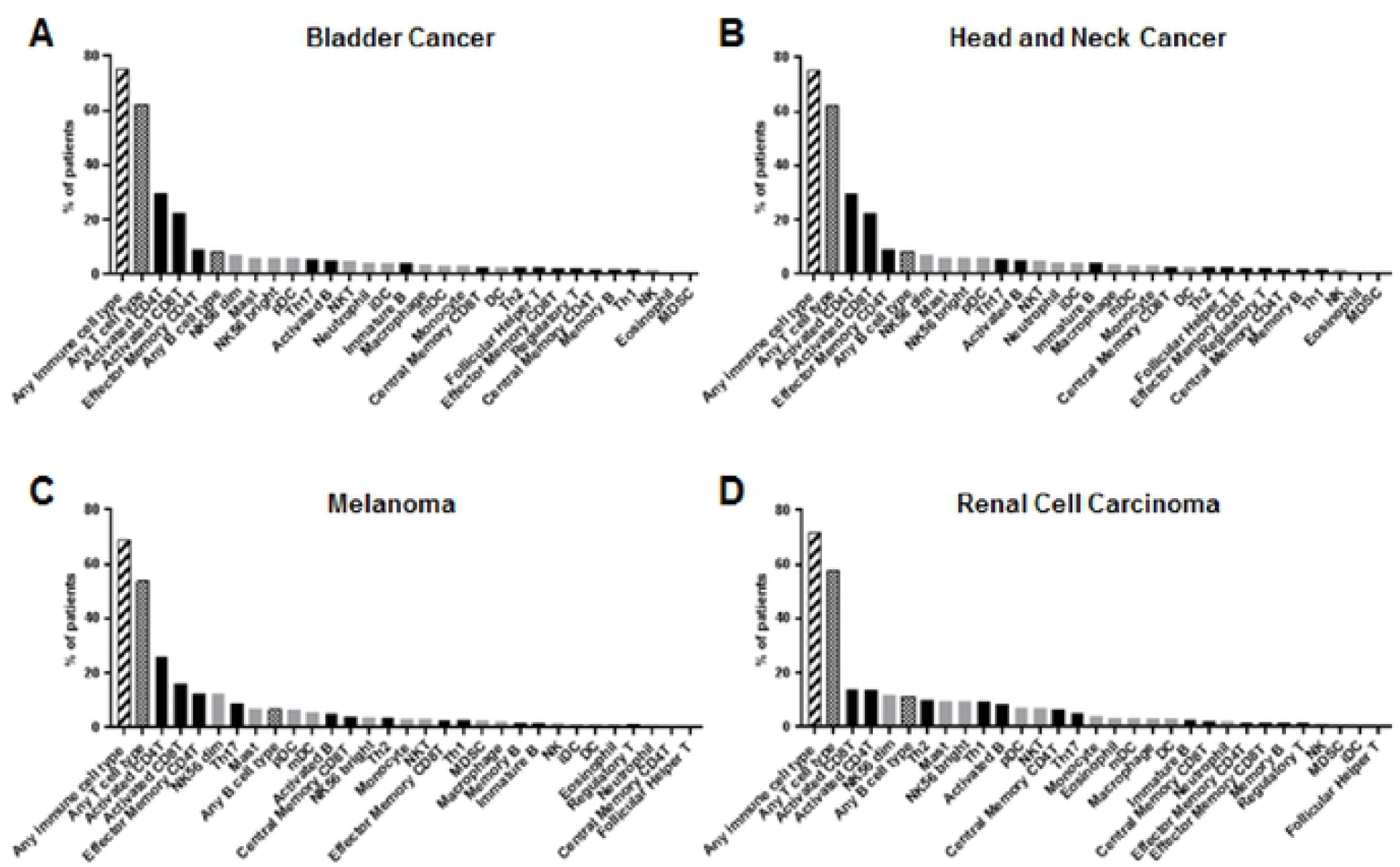
Percent of tumors infiltrated by each immune cell type in (**A**) bladder cancer, (**B**) head and neck cancer, (**C**) melanoma, and (**D**) renal cell carcinoma. NK56 bright, CD56 bright natural killer cell; DC, dendritic cell; iDC, immature dendritic cell; pDC, plasmacytoid dendritic cell; mDC, mature dendritic cell; NKT, natural killer T cells; MDSC, myeloid-derived suppressor cells (Bold bars = adaptive immune cell types, gray bars = innate immune cell types).

**Supplementary Figure 2.**
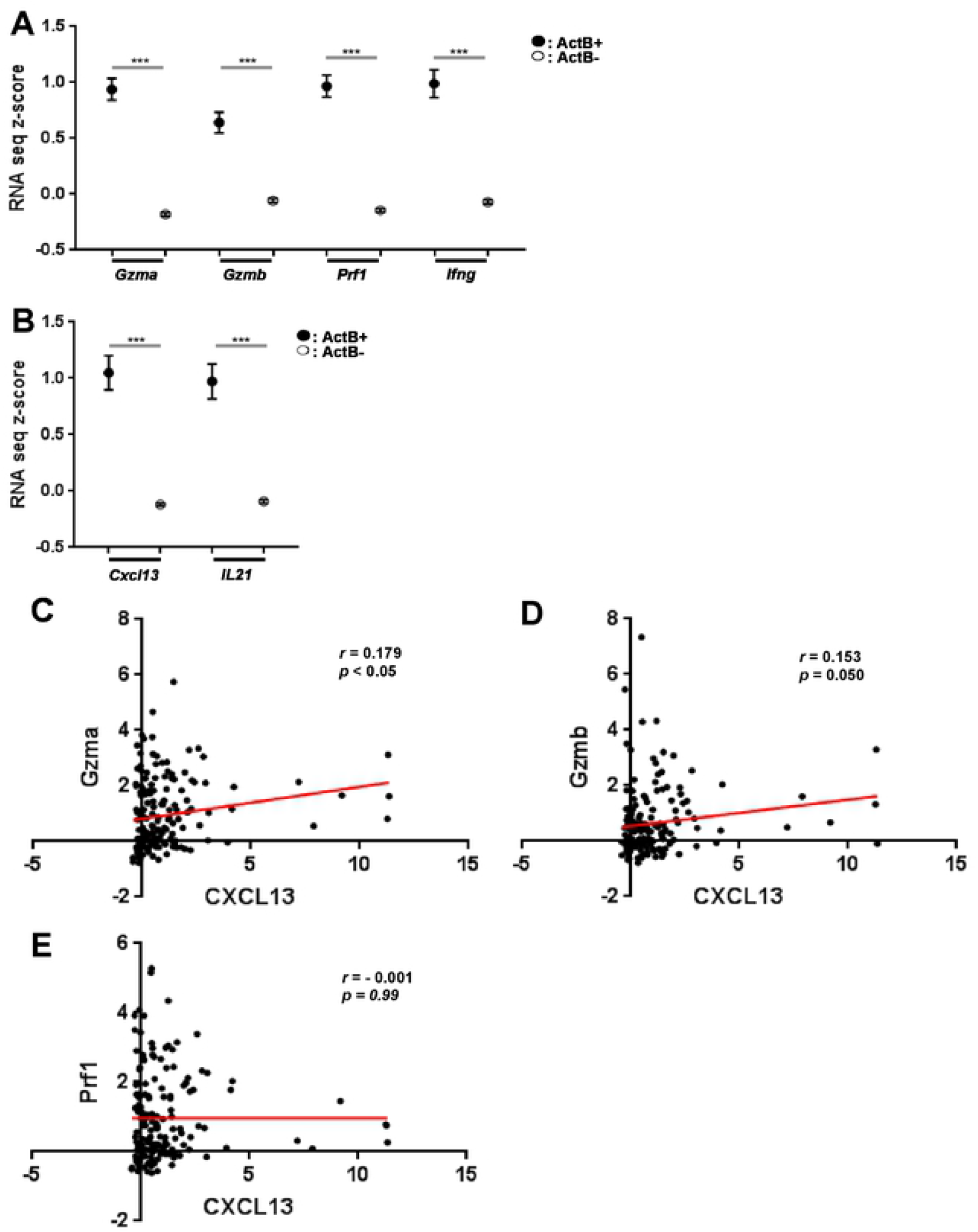
Expression scores in TLS associated markers in relation to activated B cell infiltration: (**A**) cytotoxicity-associated markers, and (**B**) TLS-associated markers *CXCL13* and *IL21*. Pearson’s coefficeints between cytotoxic effectors Granzyme A (*GzmA*), Granzyme B (*Gzmb*), Perforin (*Prf1*), and the chemokine CXCL13 in ActB+ tumors (**C-E**).

**Supplementary Figure 3.**
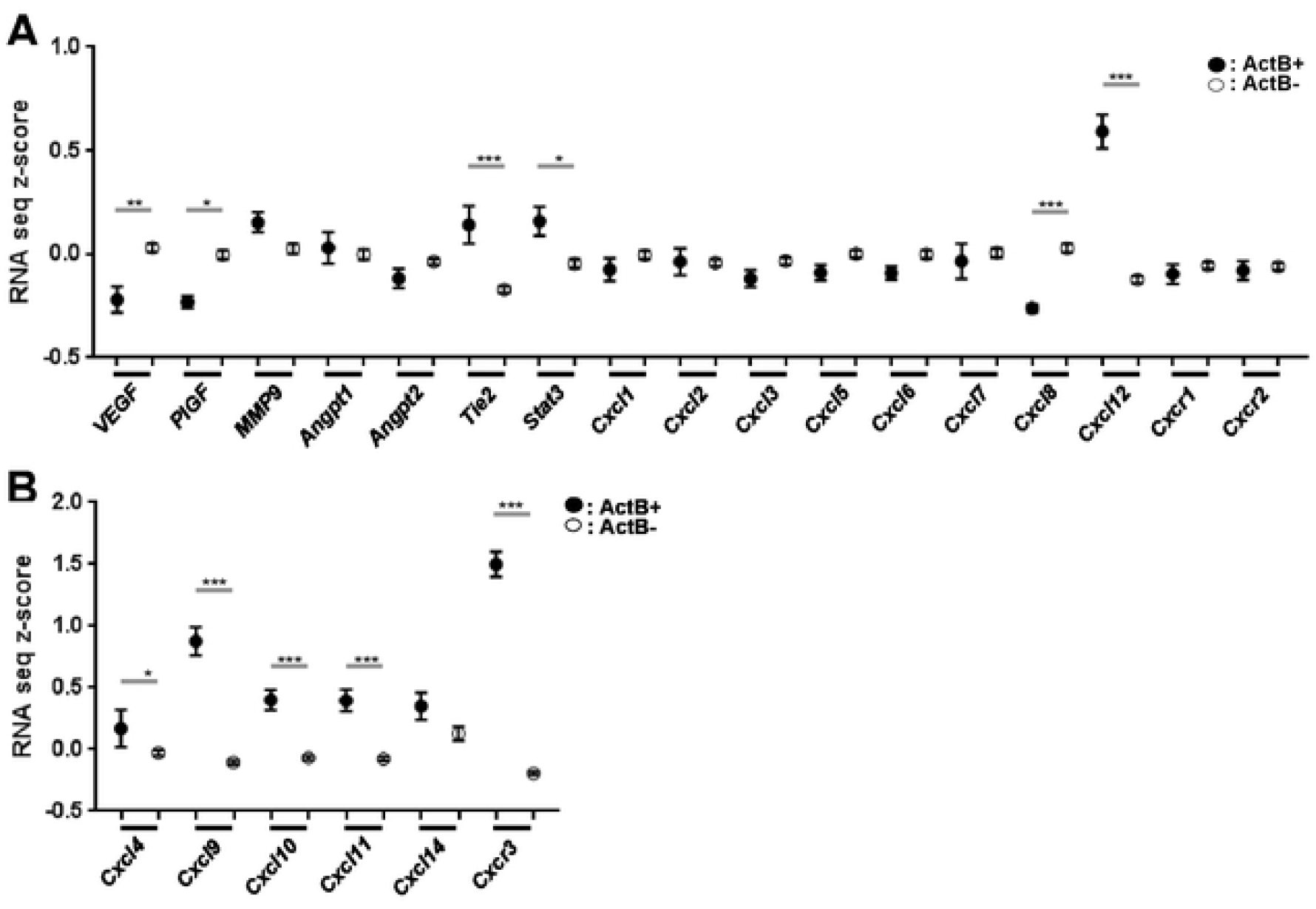
Expression levels of (**A**) angiogenic and (**B**) anti-angiogenic markers in relation to activated B cell infiltration.

**Supplementary Figure 4.**
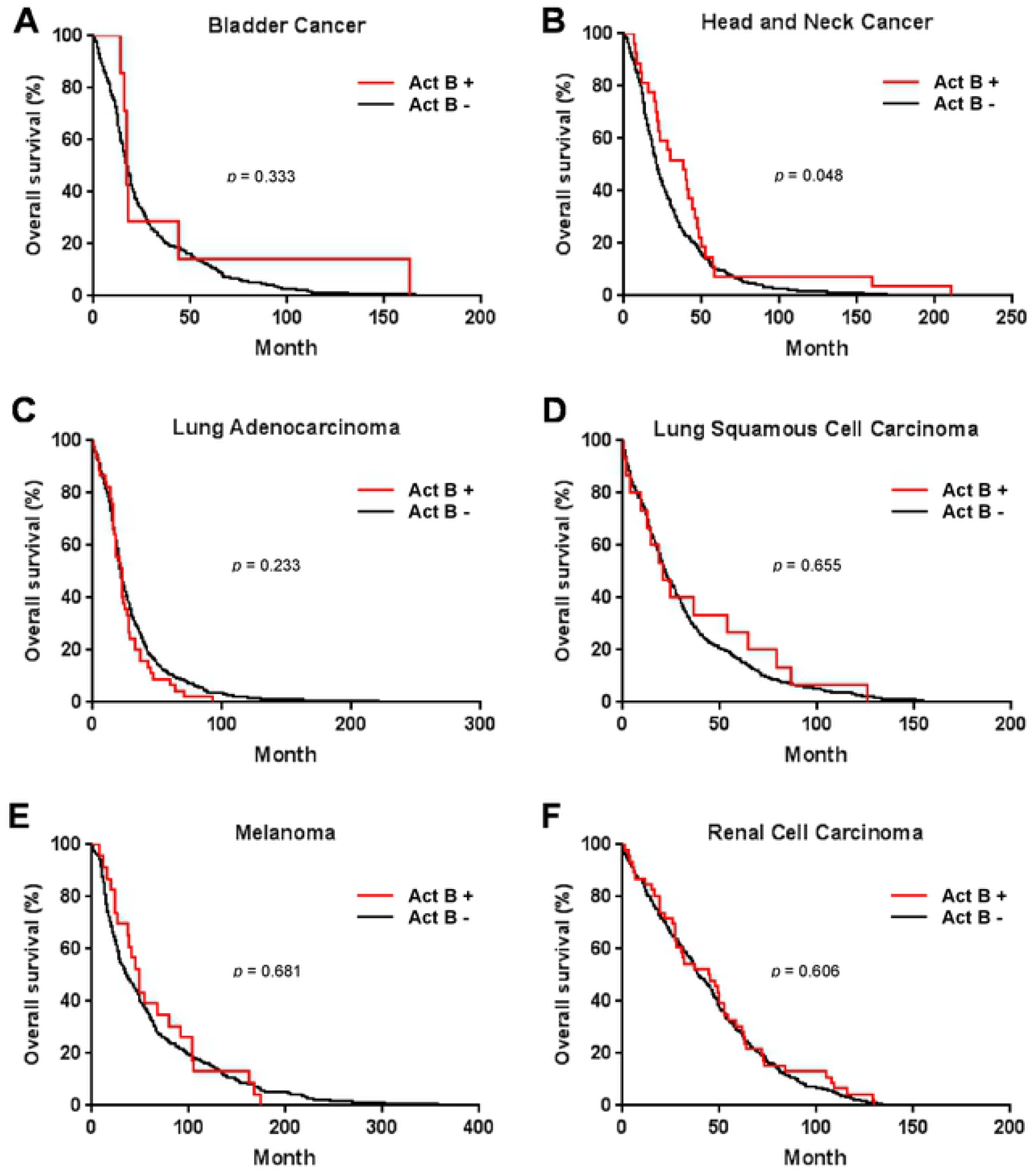
Overall survivals of patients having tumors with (red) versus without (black) activated B cell infiltration in (**A**) bladder cancer, (**B**) head and neck cancer, (**C**) lung adenocarcinoma, (**D**) lung squamous cell carcinoma, (**E**) melanoma, and(**F**) renal cell carcinoma.

**Supplementary Figure 5.**
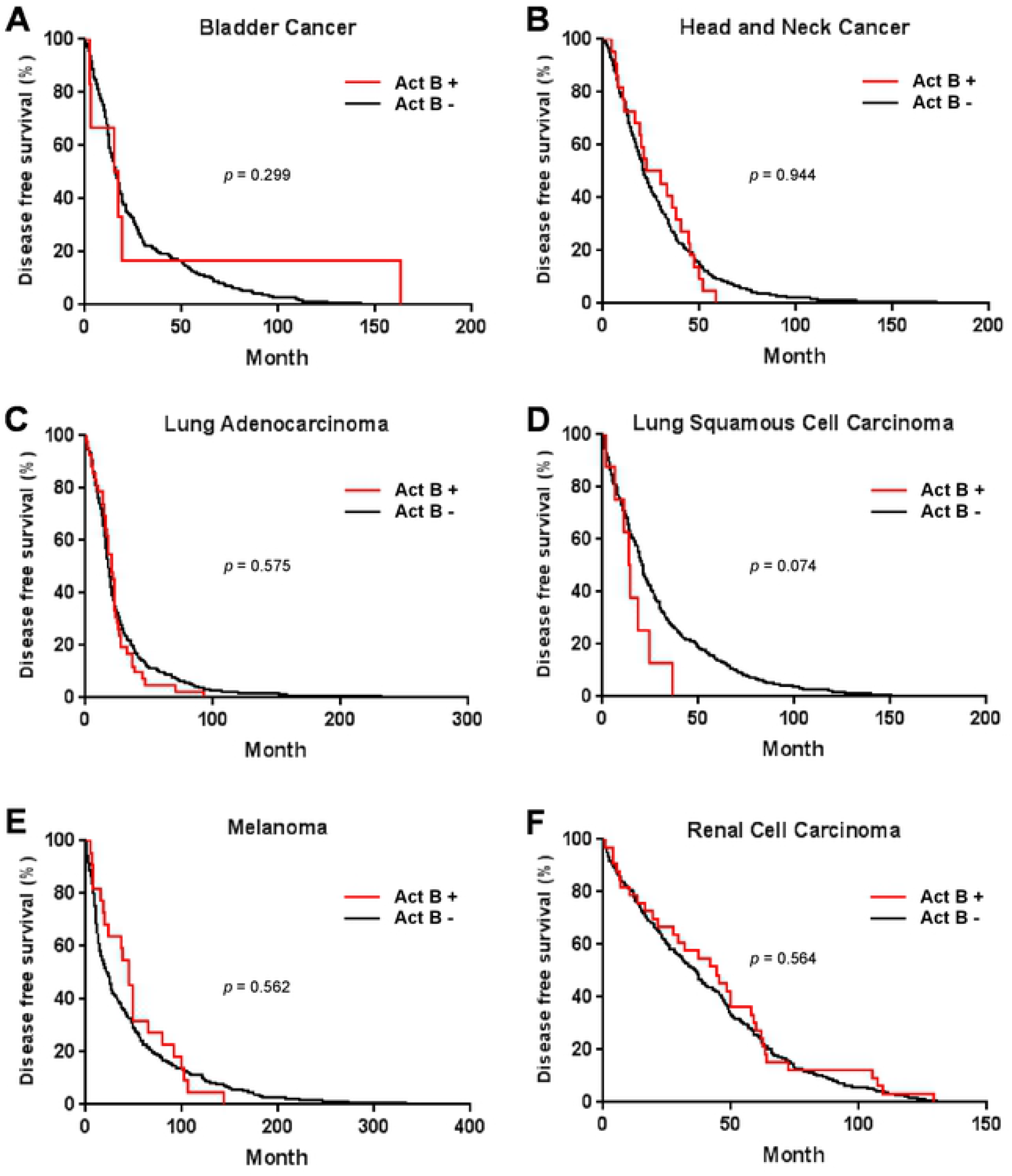
Disease free survivals of patients having tumors with (red) versus without (black) activated B cell infiltration in (**A**) bladder cancer, (**B**) head and neck cancer, (C) lung adenocarcinoma, (**D**) lung squamous cell carcinoma, (**E**) melanoma, and (**F**) renal cell carcinoma.

## Notes

The authors declare no potential conflicts of interest.

